# SHIP2 deletion in cartilage does not modulate osteoarthritis in a post-traumatic mouse model

**DOI:** 10.64898/2026.05.04.722576

**Authors:** Ana Victoria Rojo-García, Frederique M. F. Cornelis, Leire Casas-Fraile, Stéphane Schurmans, Silvia Monteagudo, Rik J. Lories

**Author notes:** These co-authors contributed equally (Ana Victoria Rojo-García, Frederique M. F. Cornelis). Corresponding author <u>Correspondence</u> Prof. Rik Lories, Laboratory of Tissue Homeostasis and Disease, Skeletal Biology and Engineering Research Centre, Department of Development and Regeneration, KU Leuven 3000 Leuven Belgium, Rik J. Lories. Ana Victoria Rojo-García. Stéphane Schurmans.

## Abstract

**Objectives:** The inositol phosphatase SHIP2 plays a crucial role in skeletal development and chondrocyte differentiation, and mutations in *INPPL1* (encoding SHIP2) cause opsismodysplasia, a chondrodysplasia with marked cartilage abnormalities. We investigated whether SHIP2 contributes to structural joint remodeling in osteoarthritis (OA).

**Methods:** A cartilage-specific conditional knockout of SHIP2 was generated using *Ship2*^fl/fl^ mice crossed with Aggrecan*Cre*^ERT2^ mice. OA was induced at 9 weeks of age via destabilization of the medial meniscus (DMM). Sham surgery served as control. Mice were sacrificed 12 weeks post-surgery. Histological evaluation of articular cartilage, synovium, osteophytes, and subchondral bone was performed. Chondrocyte hypertrophy was assessed by type X collagen (COLX) staining, and SHIP1 was evaluated as a potential compensatory mechanism.

**Results:** DMM surgery induced OA-like changes in all genotypes, including cartilage damage, synovial inflammation, osteophyte formation, and subchondral bone thickening. However, *Ship2*^cCART-KO^ mice showed no differences in OA-related parameters compared to control littermates. COLX expression increased following DMM surgery, independent of SHIP2 deletion. SHIP1 protein levels were not elevated in SHIP2-deficient mice.

**Conclusion:** These findings indicate that SHIP2, while essential for cartilage development, does not act as a structural disease modifier in post-traumatic OA, suggesting that within this context, SHIP2 is not required for maintaining adult articular cartilage structure and is unlikely to represent a major therapeutic target for modifying structural disease progression.

## INTRODUCTION

Osteoarthritis (OA) is a musculoskeletal disease characterized by articular cartilage damage^1^. It is currently the most common joint disease with an estimated 595 million people worldwide living with OA^2^. Along with cartilage degradation, OA is also associated with synovial inflammation, subchondral bone thickening, and osteophyte formation^3, 4^. It is a major cause of disability and reduced quality of life, in particular in the elderly^2^. There are no curative therapies for OA, only symptomatic treatments such as pain management and joint replacement. The complex nature of the disease and the multiple changes that occur in the joint likely contribute to the difficulty in identifying effective therapies.

In recent years, research on OA has uncovered many molecular pathways involved in disease development. However, there are still no clinically validated therapeutic targets that have resulted in approved disease-modifying drugs. As a result, there is sustained interest in identifying molecular regulators of cartilage integrity and chondrocyte behavior that could influence structural disease progression. Investigating pathways and molecules linked to other skeletal disorders such as achondroplasia or chondrodysplasias could provide novel insights into cartilage degradation and help to distinguish developmental regulators from drivers of adult joint pathology.

In 2013 homozygous or compound heterozygous mutations in *INPPL1* (inositol polyphosphate phosphatase-like 1) were reported to be the cause of opsismodysplasia, a rare chondrodysplasia characterized by growth plate defects and delayed bone maturation^5, 6^. Histological analysis of opsismodysplasia revealed increased chondrocyte density, a shortened hypertrophic chondrocyte zone, and a lack of columnar organization in the growth plates of long bones^6, 7^. These mutations frequently affect the catalytic domain of the enzyme, resulting in predicted loss of function variants. *INPPL1* encodes SHIP2 (src homology 2 domain-containing inositol phosphatase 2), a phosphatase that regulates the Akt signaling pathway and can modulate vesicular trafficking, cytoskeletal reorganization, and cell proliferation and survival, among other functions. Knockout of the catalytic domain of SHIP2 in mice causes reduced height of the hypertrophic chondrocyte zone in the growth plate and skeletal malformations^8^.

Given the essential role of SHIP2 in growth plate organization and phosphoinositide-dependent signaling, we investigated whether SHIP2 also contributes to osteoarthritis-associated joint remodeling in adult cartilage. Using a cartilage-specific, inducible deletion strategy, we examined whether loss of SHIP2 alters structural disease features in a murine post-traumatic OA model. Using comprehensive histological and molecular analyses of joint tissues, we examined cartilage damage, synovial inflammation, osteophyte formation, subchondral bone changes, and markers of chondrocyte hypertrophy. We find that SHIP2 does not influence OA progression, suggesting that its functional role may be largely confined to cartilage development rather than adult articular cartilage homeostasis.

## METHODS

### Animal model

Mice were housed in groups of 4–5 mice in a static micro-insulator cage equipped with a Macrolon filter and bedding material (composed of spruce particles of approximately 2.5–3.5mm, type Lignocel® BK 8/15), under conventional laboratory conditions (14-hour light/10-hour dark; 23 ± 2 °C), with standard mouse chow food (Sniff, Soest, Germany) and water provided *ad libitum*. All animal experiments and procedures were approved by the Ethics Committee for Animal Research at KU Leuven (P004-2022).

Cartilage-specific conditional knockout of *Ship2* (*Ship2*^cCART-KO^) was generated by crossing homozygous *Ship2*^fl/fl^, previously described^7^, and *Aggrecan*Cre^ERT2^ mice^9^. To induce *Cre*-mediated recombination, eight-week-old mice were injected intraperitoneally with tamoxifen (1 mg/mL, 200 μL/injection, three injections administered on alternate days). Male littermates homozygous for the floxed allele but lacking *Aggrecan*Cre^ERT2^, and also treated with tamoxifen, were used as controls. Genotyping was performed with PCR on genomic ear DNA, as previously described^7^.

Post-traumatic OA was induced in male control and *Ship2*^cCART-KO^ mice at 9 weeks of age using the destabilization of the medial meniscus model (DMM)^10^. Sham surgery was performed as control. The knees were harvested at 21 weeks of age (fig. 1A). Only male mice were used, consistent with standard DMM protocols.

**Figure 1.**
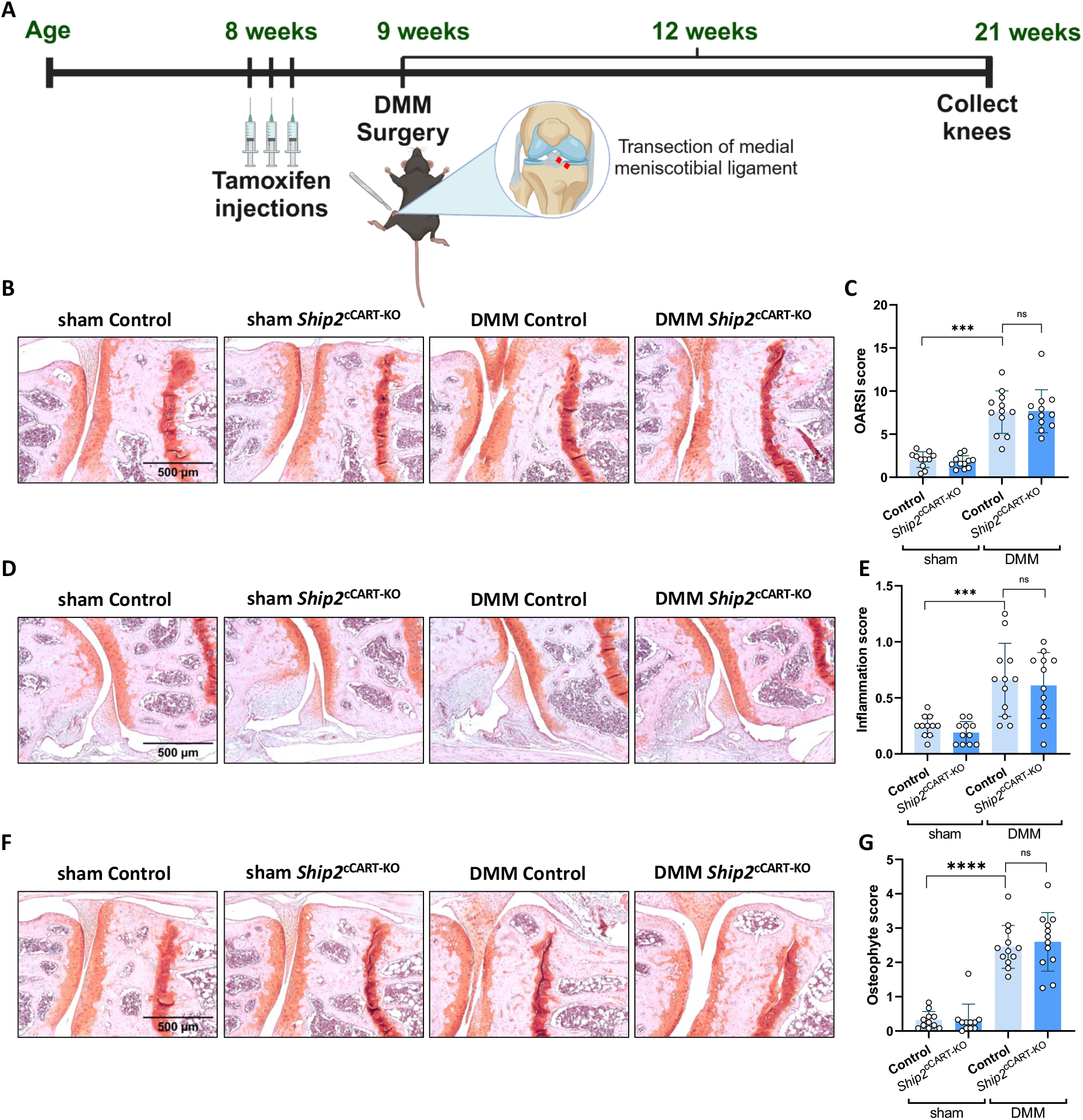
SHIP2 deletion does not affect osteoarthritic changes in cartilage, synovium, or osteophytes. **(A)** Schematic representation of the experimental design, including DMM or sham surgery in *Ship*2^cCART-KO^ and control mice. **(B, D, F)** Frontal hematoxylin-safraninO-stained sections of the lateral tibia and femur. Scale bar = 500 μm. **(C, E, G)** Quantification of cartilage damage **(C)**, synovial inflammation **(E)**, and osteophyte formation **(G)** using OARSI guidelines. (n=11 sham control n=11 sham *Ship2*^cCART-KO^, n=12 DMM control, n=12 DMM *Ship2*^cCART-KO^). Data were analyzed by two-way ANOVA after log-transformation with Tukey’s post hoc test; p-values: ns (not significant), p > 0.05, *** p < 0.001, **** p < 0.0001.

### Histology

Knees were dissected, fixed overnight at 4°C in 2% formaldehyde, and decalcified for 3 weeks in 0.5 M EDTA (pH 7.5). The knees were embedded in paraffin and cut into 5 μm sections. Severity score was determined on 6 sections spaced 100 μm apart throughout the knee and stained with hematoxylin-safranin O. Cartilage damage and synovial inflammation were measured based on OARSI guidelines^11^. The score is the sum of all four quadrants (tibia and femur, lateral and medial sides). Synovial inflammation and osteophytes were assessed using a semi-quantitative scale from 0 to 3^11, 12^. Images were acquired using a Visitron Systems microscope (Leica Microsystems GmbH).

### Subchondral bone plate histomorphometry

Histomorphometry was performed on hematoxylin-safranin O-stained sections using an Olympus DP73 microscope and digital image analysis using Osteomeasure software^13^. First, a box with a fixed width (800 μm) and variable height was drawn, with the upper limit at the transition of calcified cartilage to subchondral bone, and the lower limit located at the transition between subchondral bone and the growth plate. This zone was defined as the subchondral bone area. Next, a second box was drawn with the upper limit matched to the first box and the lower limit at the transition of the subchondral bone plate to trabecular bone. This was referred to as subchondral bone plate area. Surface areas were quantified using Osteomeasure software. To correct for sectioning artifacts, subchondral bone plate thickness was expressed as a ratio of the subchondral bone plate area to the subchondral bone area. Each mouse measurement represents the average of three sections, separated by 200 μm, and stained with hematoxylin-safranin O.

### Immunohistochemistry

Immunohistochemistry was performed on paraffin-embedded, decalcified knee sections of 5 μm thickness. For SHIP1 detection, antigen retrieval was performed with citrate-EDTA buffer (pH 6,2) for 10 min at 95°C. Next, endogenous peroxidase activity was quenched in 3% H_2_O_2_ in methanol for 10 min. Sections were blocked for 30 min in 10% goat serum in TBS-T. Primary antibody for SHIP1 (dilution: 1:50 in TBS-T, Abcam, ab45142) was applied at 4°C overnight. Rabbit IgG (Santa Cruz, sc-2027) was used as a negative control. Afterwards, sections were incubated with biotin-conjugated goat anti-rabbit IgG (Vectastain ABC kit, Vector Laboratories) for 30 min.

For the detection of Collagen type X (COLX), antigen retrieval was performed enzymatically with 10 mg/ml of Hyaluronidase (Sigma Aldrich H3884) in MgCl_2_-free PBS for 40 min at 37°C. Endogenous peroxidase activity was blocked after primary antibody incubation (2 times 10 min each). Slides were then treated with Vectastain mix (as indicated by manufacturer) for another 40 min. Lastly DAB was added. Pictures were taken using a Visitron Systems microscope (Leica Microsystems GmbH).

Quantification of DAB staining by digital image analyses after immunohistochemistry was performed in a blinded manner using the Color Deconvolution Plugin (Jacqui Ross, Auckland University) in ImageJ Software (NIH, Bethesda, USA). Quantification was based on the average of two technical replicates per mouse per condition, and staining intensity was reported % relative to the mean value of sham control mice. (100%)

### Statistics

All analyses were performed using GraphPad Prism version 8.4.3. We hypothesized that the dependent variables (OA severity, subchondral bone plate thickness, and SHIP1 and COLX protein levels) could be influenced by two independent variables (genotype and surgery). The effects of genotype, surgical intervention, and their interaction were tested using two-way analysis of variance (2-way ANOVA). The assumptions underlying ANOVA were assessed by QQ, residuals and homoscedasticity plots. For OARSI and inflammation scores data were log-transformed for better model fit. If a significant interaction was detected, a post-hoc Tukey test accounting for multiple comparisons was applied. Adjusted *P*-values < 0.05 indicated statistically significant differences. Data are reported as the differences of means (95% CI; *P*-value). Values were transformed back to report effect sizes where applicable.

## RESULTS

### Cartilage-specific conditional knockout of the SHIP2 catalytic domain does not impact OA development in the DMM model

We first investigated whether SHIP2 plays a role in the development of OA. To this end, a cartilage-specific conditional knockout of the SHIP2 catalytic domain was generated using a transgenic mouse model^7^. Male mice underwent DMM surgery, in which the medial meniscus ligament was sectioned to cause knee destabilization (fig. 1A)^10^. Knees were harvested 12 weeks post-surgery for severity scoring according to OARSI guidelines^11^. OARSI scores were 4.3-fold higher in DMM mice compared with sham operated mice (95% CI, 3.3 – 5.6, *P* < 0.0001) (fig. 1B and 1C). However, DMM *Ship2*^cCART-KO^ mice showed no difference in OARSI scores compared to DMM control mice (ratio 0.97 [95% CI, 0.6 − 1.6], *P* = 0.99).

Inflammation scores were 2.8 fold higher in DMM mice compared to sham-operated mice as expected (95% CI, 2.0 – 4.0, *P* < 0.0001), but no difference was observed between DMM *Ship2*^cCART-KO^ mice and DMM control mice (ratio 1.13 [95% CI, 0.60 − 2.14], *P* = 0.9513) (fig. 1D and 1E). As expected osteophyte formation was increased after DMM surgery (ratio 10 [95% CI, 7.1 − 15.2], *P* <0.0001) but not different between DMM *Ship2*^cCART-KO^ mice and DMM control mice (ratio 0.97 [95% CI, 0.5 − 1.9], *P* = 0.99) (fig. 1F and 1G). These data confirm that while the surgical intervention successfully induced OA-like changes, SHIP2 deletion in cartilage does not exacerbate cartilage damage, inflammation, or osteophyte formation.

### SHIP2 loss of function does not lead to changes in subchondral bone plate thickness in mice

Next, we studied modifications in the subchondral bone. Subchondral bone plate thickness was quantified as the ratio of subchondral bone plate and subchondral bone area, using hematoxylin-safranin O-stained sections (fig. 2A)^13^. A significant increase in subchondral bone plate thickness/bone area was observed in DMM mice compared to sham controls (difference of means 0.0 [95% CI, 0.001 − 0.05], *P* = 0.0414). However, no difference was found between DMM *Ship2*^cCART-KO^mice and DMM control mice (difference of means 0.013 [95% CI, −0.03 − 0.06], *P* = 0.9842) (fig. 2B). The surgical effect was most pronounced on the medial tibial side (fig. 2C) (difference of means 0.04 [95% CI, 0.006 − 0.08], *P* = 0.0238) while the lateral side remained unchanged (difference of means 0.01 [95% CI, −0.015 − 0.03], *P* = 0.4959) (fig. 2D). Thus, SHIP2 does not contribute to the bone–cartilage unit response in post-traumatic OA.

**Figure 2.**
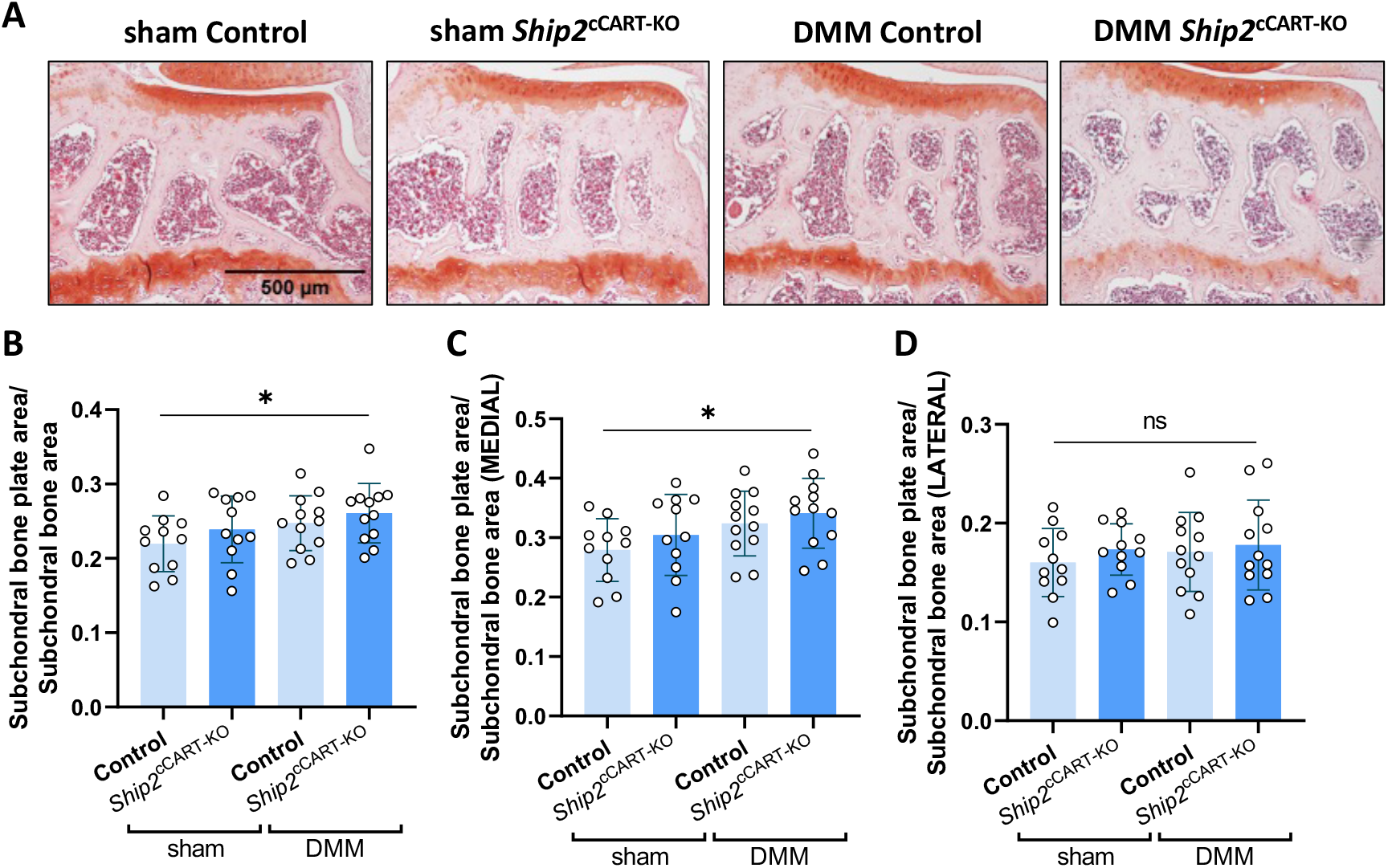
Subchondral bone plate thickness increases after DMM surgery, independent of SHIP2 inactivation. **(A)** Representative frontal hematoxylin-safranin O-stained sections of the medial tibia. Scale bar = 500 μm. **(B)** Quantification of subchondral bone plate thickness (total tibial plateau). **(C)** Medial tibia-specific quantification. **(D)** Lateral tibia-specific quantification. (n=11 sham control n=11 sham *Ship2*^cCART-KO^, n=12 DMM control, n=12 DMM *Ship2*^cCART-KO^). Data analyzed by two-way ANOVA with Tukey’s post hoc test: p-values: ns (not significant), p > 0.05, * p < 0.05.

### COLX protein expression is not altered by the knockout of SHIP2

Shortening of the hypertrophic chondrocyte zone has been described in histological samples of opsismodysplasia patients as well as animal knockouts of SHIP2 and models with pharmacological inhibition of this enzyme^6^. In OA, chondrocytes undergo a change from a quiescent phenotype towards a hypertrophic profile^1^. To investigate whether SHIP2 deletion influences hypertrophic differentiation in articular cartilage, immunostaining for COLX, a known marker of chondrocyte hypertrophy^14^ was performed (fig. 3A). We observed a 21.9 fold increase of COLX in DMM mice compared to sham-operated animals (95% CI, 11.2 – 42.9, *P* < 0.0001) (fig. 3B). However, there was no difference between DMM control mice and DMM *Ship2*^cCART-KO^ mice (ratio 0.87 [95% CI, 0.28 – 2.69], *P* = 0.983), suggesting that SHIP2 knockout does not modulate chondrocyte hypertrophy in this context.

**Figure 3.**
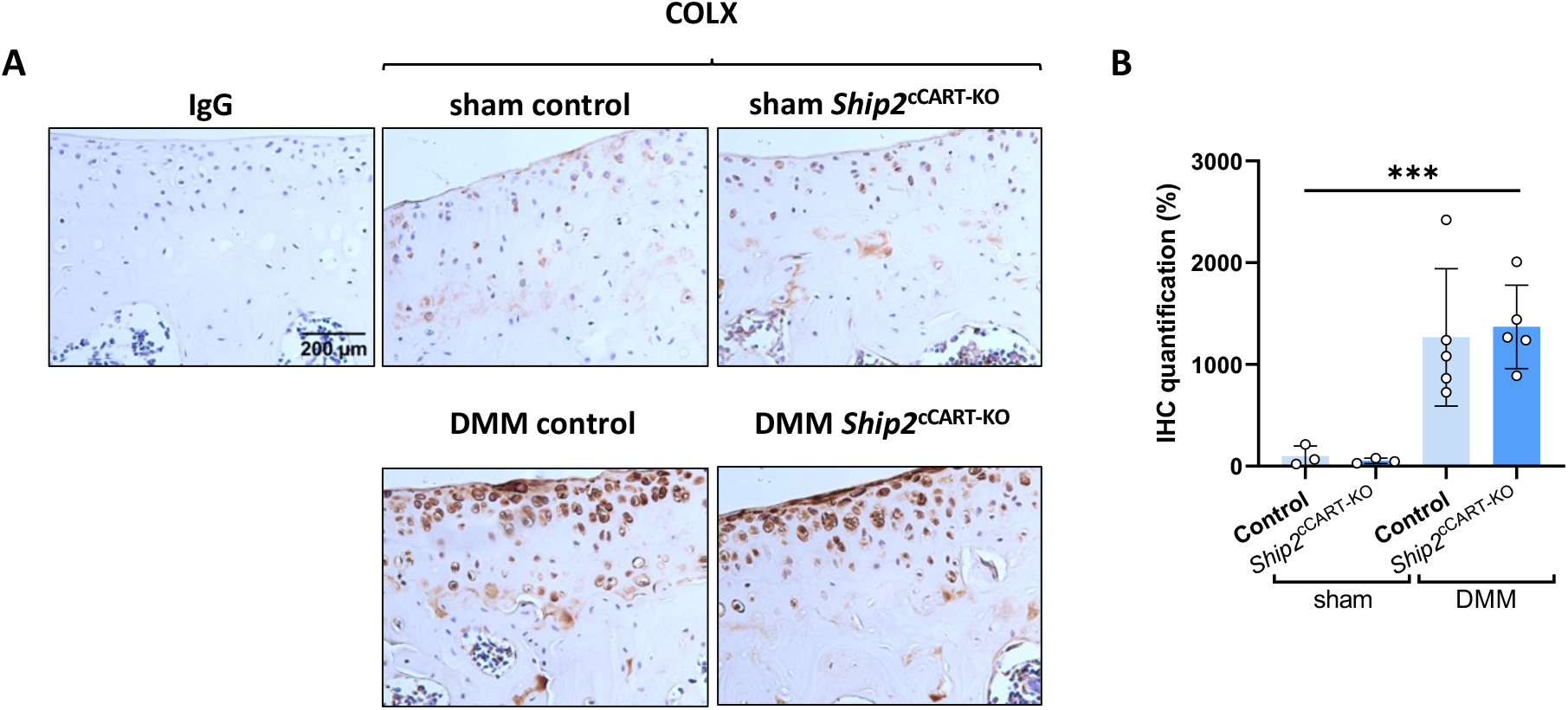
COLX expression increases after DMM surgery but is not altered by SHIP2 inactivation. **(A)** Immunohistochemical detection of type X collagen (COLX) in the articular cartilage of sham control, sham *Ship2*^cCART-KO^, DMM control, and DMM *Ship2*^cCART-KO^ mice. Scale bar = 200 μm.**(B)** Quantification of COLX staining intensity. (n=3 sham control, n=3 sham *Ship2*^cCART-KO^, n=5 DMM control, n=5 DMM *Ship2*^cCART-KO^). Data analyzed by two-way ANOVA after log-transformaiton with Tukey’s post hoc test: p-values: ns (not significant), p > 0.05.

### SHIP1 is not upregulated upon the knockout of SHIP2 or DMM intervention

We next examined whether SHIP1, a paralog of SHIP2^15-17^, is upregulated as a compensatory mechanism in response to SHIP2 deletion in *Ship2*^cCART-KO^ mice. While SHIP2 is ubiquitous, SHIP1 is mostly expressed in hematopoietic lineage and bone-forming lineage cells^15, 16, 18^. Both phosphatases modulate the PI3K-Akt pathway^19^. Immunohistochemical staining of SHIP1 revealed slightly reduced expression in DMM compared to sham-operated mice (fig. 4A) (ratio 0.82, [95% CI, 1.00 – 1.49], *P* = 0.0496), but no differences were detected between *Ship2*^cCART-KO^ and control mice (ratio 1.10 [95% CI, 0.79 − 1.53], *P* = 0.83) (fig. 4B). These findings indicate that SHIP1 is not upregulated upon SHIP2 deletion or DMM intervention, and its expression is therefore not a compensatory factor in the model used.

**Figure 4.**
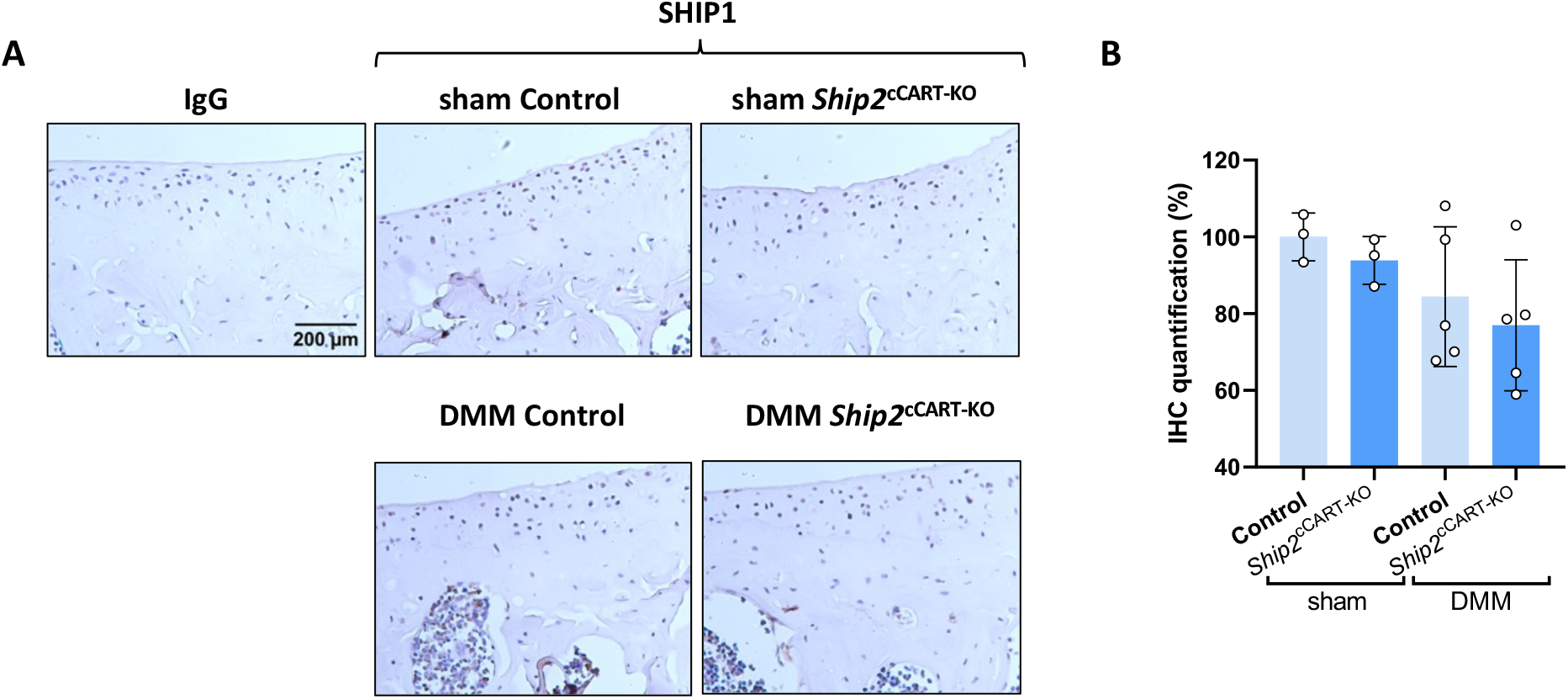
SHIP1 protein levels are not increased in *Ship2*^cCART-KO^ mice. **(A)** Immunohistochemical staining of SHIP1 in articular cartilage from sham control, sham *Ship2*^cCART-KO^, DMM control, and DMM *Ship2*^cCART-KO^ mice. Scale bar = 200 μm. **(B)** Quantification of SHIP1 staining intensity. (n=3 sham control, n=3 sham *Ship2*^cCART-KO^, n=5 DMM control, n=5 DMM *Ship2*^cCART-KO^). Data analyzed by two-way ANOVA after log-transformation with Tukey’s post hoc test: p-values: ns (not significant), p > 0.05.

## DISCUSSION

Understanding which molecular pathways actively drive cartilage degradation in OA remains a major challenge. While numerous signaling pathways are involved in skeletal development and chondrocyte differentiation, it is increasingly clear that not all of these developmental regulators have roles in adult joint disease. In this study, we investigated whether SHIP2, a phosphoinositide phosphatase with a well-established role in growth plate organization and skeletal development, contributes to structural joint changes in a murine model of post-traumatic OA. Using a cartilage-specific, inducible deletion strategy in adult mice, we show that loss of SHIP2 does not influence cartilage damage, synovial inflammation, osteophyte formation, subchondral bone remodeling, or chondrocyte hypertrophy following destabilization of the medial meniscus.

SHIP2 is essential for normal endochondral ossification and chondrocyte differentiation during skeletal development. Mutations in SHIP2 cause opsismodysplasia, a severe chondrodysplasia characterized by growth plate disorganization, shortened hypertrophic zones, and impaired bone maturation^5, 6, 20, 21^. Consistent with these human phenotypes, catalytic inactivation of SHIP2 in mice leads to pronounced developmental skeletal abnormalities^8^. These observations provided a strong rationale to examine whether SHIP2 might similarly regulate chondrocyte responses under pathological conditions in the adult joint, where OA is associated with phenotypic changes that partially resemble aspects of chondrocyte hypertrophy and developmental reprogramming.

Despite this rationale, cartilage-specific deletion of SHIP2 in adult mice had no detectable impact on disease severity in the DMM model. Histological assessment demonstrated the expected OA-associated changes induced by joint destabilization, including cartilage erosion, osteophyte formation, synovial inflammation, and subchondral bone plate thickening. However, all of these features were entirely driven by surgical intervention and occurred independently of SHIP2 expression in cartilage. These findings indicate that SHIP2 appears dispensable for the structural manifestations of post-traumatic OA once articular cartilage has reached full differentiation.

One potential explanation for this lack of effect is that SHIP2 functions primarily during developmental stages, when chondrocytes undergo proliferation, hypertrophic differentiation, and matrix remodeling as part of endochondral bone formation. In contrast, articular chondrocytes in adult cartilage are largely quiescent under physiological conditions, and their pathological activation during OA may rely on signaling pathways distinct from those governing developmental cartilage biology. Our data support the notion that reactivation of hypertrophic markers during OA does not necessarily imply reuse of all the same upstream regulatory machinery that operates during skeletal development.

The subchondral bone lies underneath the articular cartilage and provides nutrients and mechanical support^22, 23^. In OA, as cartilage degrades it exposes the underlying bone provoking abnormal growth and what has been characterized as subchondral bone plate thickening^4, 24^. Subchondral bone thickened in mice after DMM surgery but was independent of the genotype. DMM surgery replicated all OA histological traits, nonetheless the catalytic inactivation of SHIP2 did not have any effect on the subchondral bone plate. Hence, it seems that SHIP2 alters bone formation in embryonic stages and not in fully differentiated cells in adult mice.

Consistent with this interpretation, we observed a robust increase in type X collagen expression following DMM surgery, reflecting chondrocyte hypertrophy associated with OA progression^14^. However, SHIP2 deletion did not modulate COLX levels, indicating that hypertrophic differentiation in this context proceeds independently of SHIP2-mediated phosphoinositide signaling. This further suggests that the molecular control of chondrocyte hypertrophy in osteoarthritis diverges from that operating in the growth plate.

We also explored whether compensatory upregulation of the related phosphatase SHIP1 could mask a potential role of SHIP2 in adult cartilage. SHIP1 is known to regulate PI3K–Akt signaling and has been implicated in skeletal biology^16, 19^, although its expression is more restricted to hematopoietic and bone-forming lineages^16, 18^. No increase in SHIP1 protein levels was detected in SHIP2-deficient cartilage, arguing against functional compensation at the level of protein expression in this model. While this does not exclude compensatory signaling via other phosphatases or pathways, it reinforces the conclusion that SHIP2 itself is not a critical determinant of OA-associated cartilage pathology.

The skeletal phenotype of SHIP2 knockout mice is milder than in humans with SHIP2 mutations^7, 8^. Hence, the role of SHIP2 in mice may differ from its role in humans^7, 8^. Another limitation of the present study is that it focuses on a single murine model of post-traumatic OA. It therefore remains possible that SHIP2 could influence other OA mechanistic context, such as age-related or metabolic disease, or contribute to joint pathology through non-cartilaginous tissues.

In conclusion, our study shows that although SHIP2 is a critical regulator of chondrocyte differentiation and skeletal development, it does not contribute to structural joint degeneration in a murine post-traumatic osteoarthritis model when deleted in adult cartilage. These results highlight an important distinction between developmental regulators and drivers of adult joint disease and underscore the need for careful functional validation of putative OA targets in appropriate disease contexts. While SHIP2 may represent a relevant therapeutic target for developmental skeletal disorders, it is unlikely to play a major role in modifying structural disease progression in osteoarthritis.

## ACKNOWLEDGEMENTS

The authors thank Lies Storms and Ann Hens for excellent technical support.

## AUTHOR CONTRIBUTIONS

**A. V. Rojo-Garcia**: analysis and interpretation of data; drafting the manuscript and final approval of the version to be published; agreement to be accountable for all aspects of the work. **F. M. F. Cornelis**: acquisition, analysis and interpretation of data; drafting the manuscript and final approval of the version to be published; agreement to be accountable for all aspects of the work. **L. Casas-Fraile**: interpretation of the data; critical reviewing for important intellectual content; final approval of the version to be published. **S. Schurmans**: concept of the work; interpretation of the data; critical reviewing for important intellectual content; final approval of the version to be published. **S. Monteagudo**: concept of the work; interpretation of the data; critical reviewing for important intellectual content; final approval of the version to be published. **R. J. Lories**: concept of the work; interpretation of the data; critical reviewing for important intellectual content; final approval of the version to be published; agreement to be accountable for all aspects of the work.

